# Androgen receptor determines skeletal muscle sexual dimorphism

**DOI:** 10.64898/2026.08.17.745371

**Authors:** Hiroshi Sakai, Yuta Yanagihara, Kaori Tanaka, Ayaka Tabuchi, Hiroyuki Iwamoto, Yusuke Horita, Satoru Otowa, Tomofumi Kinoshita, Kunihiko Watamori, Kazunori Hino, Masaki Takao, Pascal Maire, Shahragim Tajbakhsh, Hidetaka Kosako, Tatsuya Sawasaki, Takashi Yamada, Yutaka Kano, Akihito Harada, Yasuyuki Ohkawa, Yuuki Imai

## Abstract

The molecular and functional bases of sexual dimorphism in skeletal muscle remain poorly understood. The androgen receptor (AR) is a major regulator of sex-biased gene expression in muscle, but its genomic targets and associated coregulators *in vivo* are incompletely defined. Using ChIL-seq and an AirID-AR knock-in mouse, we mapped AR-bound genes and AR-associated proteins in skeletal muscle and identified histone deacetylase-linked corepressors. We further identified myosin binding protein H (Mybph) as a female-biased AR-repressed gene conserved in mouse and human muscle. *Mybph* loss disrupted sarcomeric organization and selectively delayed postinjury force recovery in female mice. These findings define an *in vivo* AR regulatory network and identify AR-dependent *Mybph* repression as a potential mechanism contributing to skeletal muscle sexual dimorphism.

## Main text

Skeletal muscle exhibits pronounced sexual dimorphism at anatomical, functional, and molecular levels ^1^. In addition to readily apparent differences in muscle mass and morphology, several aspects of muscle function differ between the sexes ^2^; for example, females show faster recovery of muscle force than males following muscle injury or damaging eccentric contractions ^3–5^. Sexual dimorphism in skeletal muscle is also evident in its transcriptome, as the expression profiles of autosomal chromosome genes alone can distinguish male from female muscle with high accuracy ^6^. Analyses aimed at identifying the transcriptional regulators underlying these sex-biased expression patterns have ranked the androgen receptor (AR) as the most prominent hormone-responsive transcription factor^6^.

AR is a ligand-activated transcription factor that mediates the actions of androgens ^7^. In skeletal muscle, AR is expressed in muscle stem cells, mesenchymal progenitors, and differentiated myofibers ^8^. In myofibers, AR regulates genes involved primarily in muscle metabolism and contraction ^9–11^. AR is thought to control gene expression through interactions with transcriptional coregulators ^12^. However, a comprehensive understanding of the genomic targets of AR and its associated coregulators in skeletal muscle is still lacking. In particular, the AR target genes and gene regulatory mechanisms that give rise to sexual dimorphism remain poorly understood in mouse skeletal muscle *in vivo* and in human skeletal muscle.

### ChIL-seq maps AR genomic occupancy in male skeletal muscle tissue sections

To identify genomic targets of AR in skeletal muscle, we performed chromatin integration labeling followed by sequencing (ChIL-seq), a low-input method that combines immunostaining, transposase-mediated tagging, and linear amplification to profile epigenomic features and protein–DNA interactions, even from single tissue sections ^13,14^. We collected biceps brachii muscles from the forelimbs of male mice 2 h after treatment with or without dihydrotestosterone (DHT) (Fig. 1A). AR ChIL-seq detected 1,180 AR peaks in male mouse skeletal muscle. Although the signal-to-noise ratio was low in untreated samples compared with the secondary-antibody-only control, DHT treatment markedly increased the signal-to-noise ratio (Fig. 1B). These peaks were distributed across promoter, intronic, and intergenic regions (Fig. 1C). The most enriched transcription factor–binding motif within these peaks was the androgen response element (ARE), the canonical AR-binding motif (Fig. 1D), indicating that AR ChIL-seq performed as expected. To examine skeletal muscle– specific AR-binding events, we compared the AR ChIL-seq profile in skeletal muscle with conventional AR ChIP-seq profiles from prostate, kidney, and epididymis ^15^. Correlation analysis revealed a reproducible skeletal muscle–specific AR-binding pattern (Fig. 1E). Genes associated with AR peaks in skeletal muscle were enriched for Gene Ontology (GO) Biological Process terms related to “muscle structure development” and “actin filament–based processes” (Fig. 1F), consistent with previous studies of AR knockout in skeletal muscle fibers ^9,16^.

**Fig 1.**
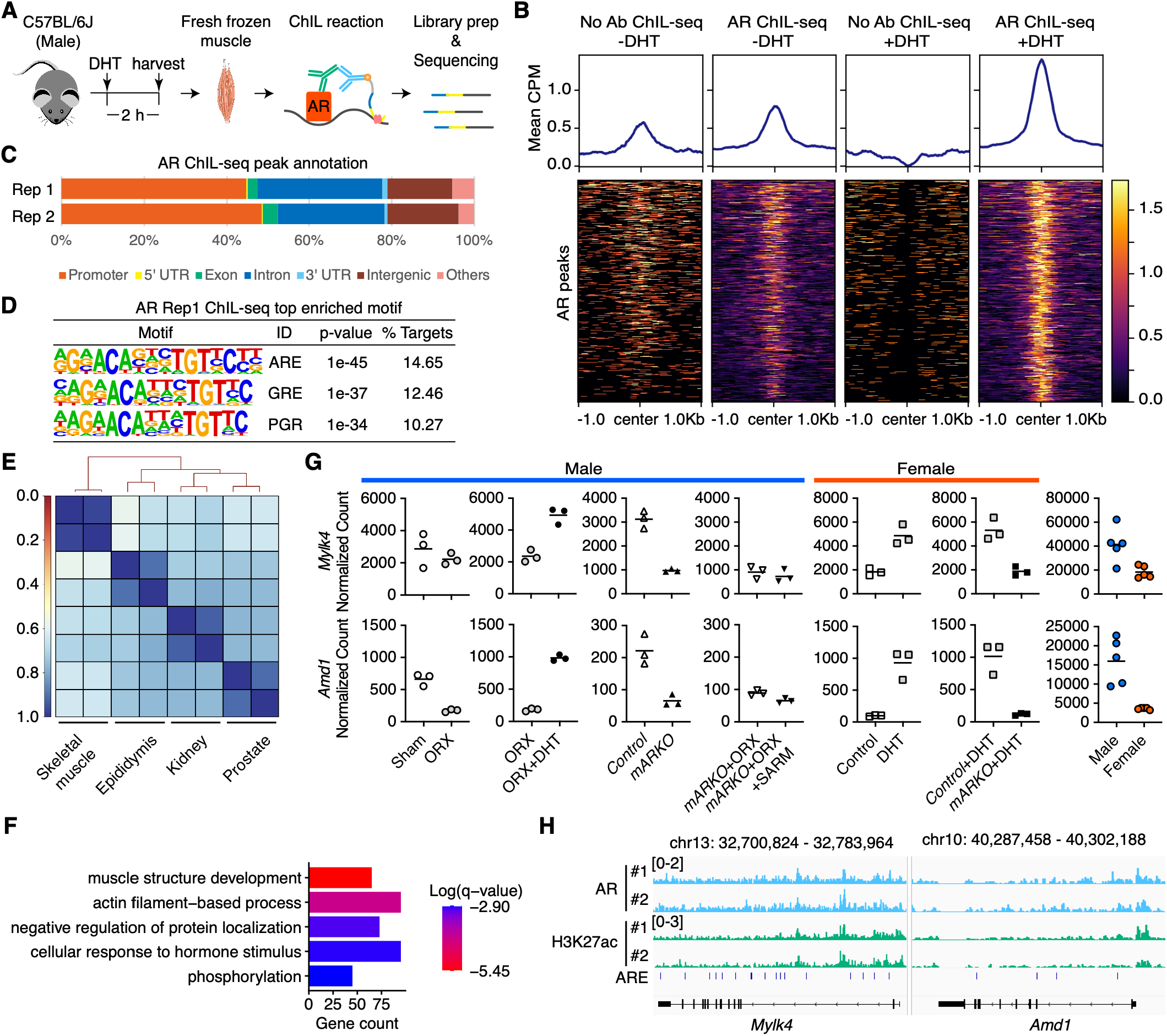
ChIL-seq analysis of AR genomic occupancy in male skeletal muscle tissue sections. (A) Schematic of ChIL-seq analysis in mouse skeletal muscle. DHT, dihydrotestosterone. (B) Aggregation plot of AR ChIL-seq signals with or without DHT treatment, centered on AR ChIL-seq peaks identified after DHT treatment. CPM, counts per million. (C) Genomic distribution of AR peaks detected by ChIL-seq in skeletal muscle. (D) Top enriched motifs identified by HOMER in AR ChIL-seq peaks after DHT treatment. ARE, androgen response element; GRE, glucocorticoid response element; PGR, progesterone receptor motif. (E) Hierarchical clustering based on Pearson correlation coefficients of AR ChIL-seq data from skeletal muscle and AR ChIP-seq data from epididymis, kidney, and prostate (GSE47192). (F) Top GO Biological Process terms associated with genes near AR ChIL-seq peaks identified after DHT treatment. (G) RNA-seq analysis of selected androgen-induced and AR-dependent genes in skeletal muscle fibers from male and female mice. Sham, sham operation; ORX, orchiectomy; Control, AR flox mice; mARKO, skeletal muscle-specific AR knockout mice; SARM, selective androgen receptor modulator. (H) Representative AR peaks after DHT treatment, together with H3K27ac ChIL-seq signals, at androgen-induced and AR-target genes in skeletal muscle fibers.

To identify androgen-responsive chromatin regions, we performed H3K27ac ChIL-seq in the androgen treatment model. We identified 2,276 regions with increased H3K27ac signals after DHT treatment (fig. S1A). These DHT-responsive H3K27ac regions were mainly distributed in intronic and intergenic regions (fig. S1B). Motif enrichment analysis showed that the ARE was the most enriched motif within these regions (fig. S1C), suggesting that AR recruitment contributes to androgen-induced chromatin activation. GO Biological Process analysis of the 1,585 genes associated with DHT-activated H3K27ac peaks showed enrichment for “actin filament–based processes” (fig. S1D), consistent with DHT–AR regulation of skeletal muscle fibers.

To assess the effects of AR binding on gene expression, we integrated ChIL-seq data with RNA-seq data from male mice under four conditions: orchiectomy (ORX), ORX plus DHT treatment, skeletal muscle fiber–specific AR knockout, and the combination of these conditions, as well as from female mice under two conditions: DHT treatment and skeletal muscle fiber–specific AR knockout ^10,11,17^. We confirmed that previously reported AR-regulated genes such as *Mylk4* and *Amd1*, were regulated by androgen through AR in skeletal muscle fibers and showed male-biased expression (Fig. 1G). These androgen-induced, skeletal muscle fiber AR–dependent genes were associated with AR peaks and DHT-induced H3K27ac sites containing AREs, as detected by ChIL-seq (Fig. 1H). We also identified *Smtnl2*, which is involved in actin cytoskeleton organization, as well as the zinc transporter *Slc30a2* and the sodium-coupled neutral amino acid transporter *Slc38a4*, as genes robustly induced by androgen–AR signaling in skeletal muscle fibers (fig. S2, A and B). Together, integration of the ChIL-seq and RNA-seq datasets not only recapitulated previously reported androgen-induced AR target genes but also uncovered previously unrecognized direct targets in skeletal muscle fibers.

### AirID-AR mice identify AR coregulators in skeletal muscle *in vivo*

AR functions with coregulators to orchestrate gene expression in a cell type–specific manner ^12^. To identify AR coregulators in skeletal muscle *in vivo*, we generated knock-in mice expressing AR fused to ancestral BirA for proximity-dependent biotin identification (AirID) (Fig. 2A and fig. S3, A to C) ^18^. AirID-AR did not affect body weight, skeletal muscle mass, including the highly androgen-sensitive levator ani/bulbocavernosus (LA/BC) muscles, or AR expression in skeletal muscle (fig. S3, D to F). To identify proteins biotinylated by AirID-AR, we performed liquid chromatography–tandem mass spectrometry (LC-MS/MS) analysis of biotinylated peptides isolated from tibialis anterior (TA) muscles of control (*Ar*^*+/Y*^) and AirID-AR (*Ar*^*AirID/Y*^) mice treated with biotin for 3 days, with or without DHT treatment (Fig. 2B). AR was the most enriched biotinylated protein in AirID-AR mice under both untreated and DHT-treated conditions, indicating that AirID-AR functioned as expected *in vivo* (Fig. 2C). We also detected several biotinylated lysine residues within an AR N-terminal region previously implicated in ligand-dependent N/C interactions and receptor dimerization (Fig. 2D) ^19,20^. In addition, the biotinylated proteins included known AR-binding proteins (Fig. 2E). GO Cellular Component analysis of proteins detected under common or DHT-induced conditions (Fig. 2F) showed enrichment for transcription regulator complexes (Fig. 2G). Notably, histone deacetylase complexes, including SIN3A and NCOR2, were also enriched (Fig. 2H). By contrast, LSD1, a histone demethylase previously reported to function as an AR-associated coregulators ^21,22^, was not detected as an AR-binding protein in this system. These findings suggest that AR may repress, as well as activate, gene expression in skeletal muscle through histone deacetylase-associated corepressor complexes. We further detected biotinylated lysine residues in NCOR2 within nuclear receptor–interaction sites (fig. S3G) ^23,24^ and confirmed coexpression of NCOR2 and AR in skeletal muscle fibers (Fig. 2I). Together, these AirID-AR mice identified AR coregulators, including corepressors, in skeletal muscle *in vivo*.

**Fig 2.**
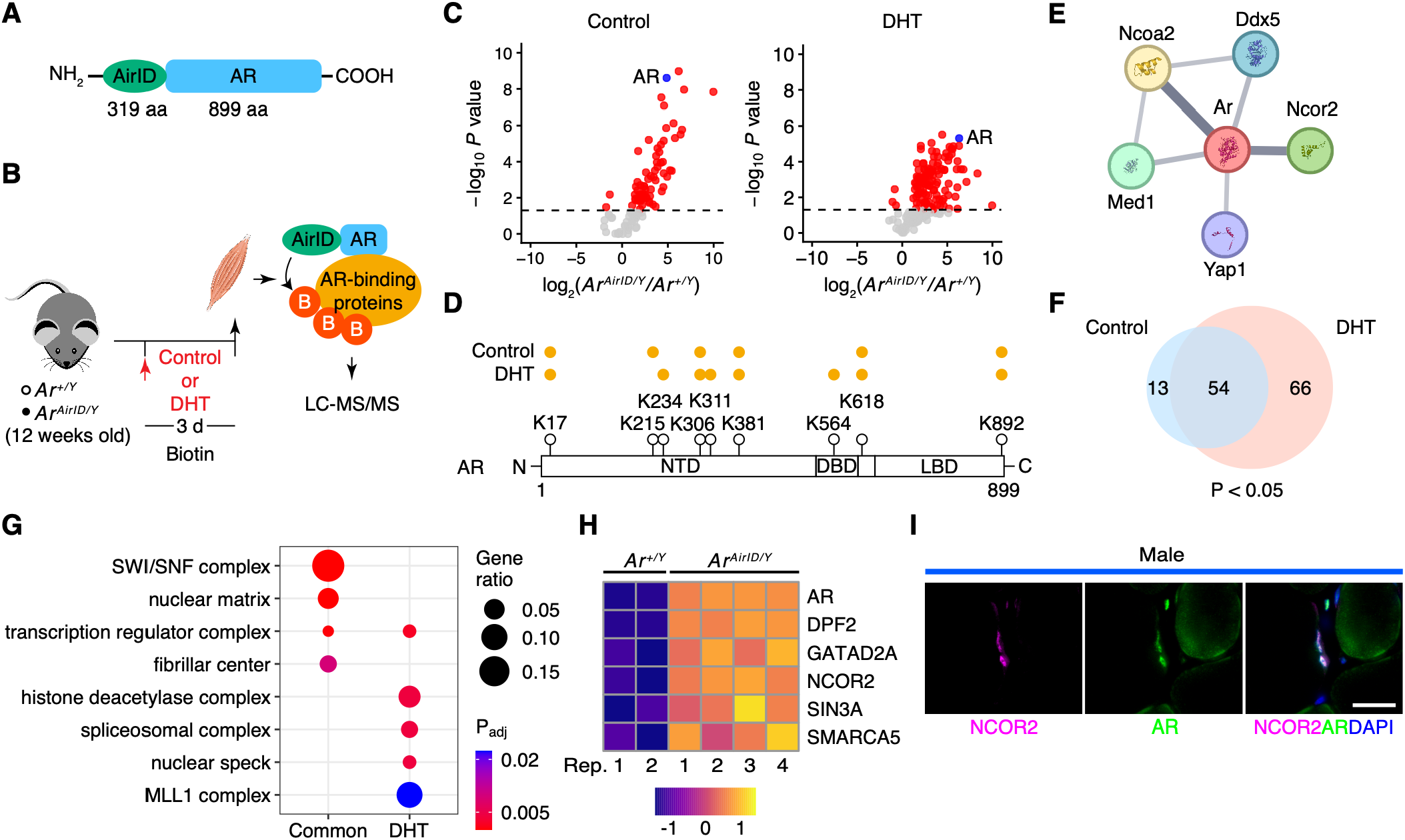
Identification of AR coregulators in skeletal muscle in vivo using AirID-AR mice. (A) Schematic of the AR protein fused to AirID. (B) Schematic of LC-MS/MS analysis to detect biotinylated proteins isolated from TA muscles of AirID-AR mice. DHT, dihydrotestosterone. (C) Volcano plots of biotinylated proteins from untreated control mice (left) and DHT-treated mice (right). Blue dots indicate AR. (D) Biotinylated sites detected in AR from AirID-AR mice with or without DHT treatment. Orange dots denote biotinylated lysine (K) residues. NTD, N-terminal domain; DBD, DNA-binding domain; LBD, ligand-binding domain. (E) Known interactions between AR and proteins detected in DHT-treated AirID-AR mice, analyzed using STRING. (F) Venn diagram of biotinylated proteins detected in skeletal muscle from untreated control and DHT-treated AirID-AR mice. (G) GO Cellular Component enrichment analysis of cellular components for proteins commonly detected under both conditions or specifically detected after DHT treatment in skeletal muscle from AirID-AR mice. (H) Heat map of detected histone deacetylase complex–related proteins in DHT-treated AirID-AR mice. (I) Immunostaining for AR and NCOR2 in male skeletal muscle. Scale bar, 20 µm.

### Loss of the sex-biased AR target Mybph impairs myofibrillar alignment and muscle force recovery in female mice

Because AirID-AR identified histone deacetylase complexes among AR-associated proteins, we searched multiple RNA-seq datasets for genes repressed by androgen in a skeletal muscle fiber AR–dependent manner. This analysis identified myosin binding protein H (Mybph), a member of the myosin-binding protein family, as an AR-repressed gene in skeletal muscle (Fig. 3A). We confirmed that *Mybph* transcripts were up-regulated after ORX in male mice (Fig. 3B). Mybph expression was also female-biased at both the transcript and protein levels (Fig. 3, C and D). AR ChIL-seq revealed an AR peak at the *Mybph* locus containing an ARE, which coincided with low H3K27ac signals in male muscle and high H3K27ac signals in female muscle (Fig. 3E). These data suggest that Mybph is a sex-biased AR direct target gene in skeletal muscle.

**Fig 3.**
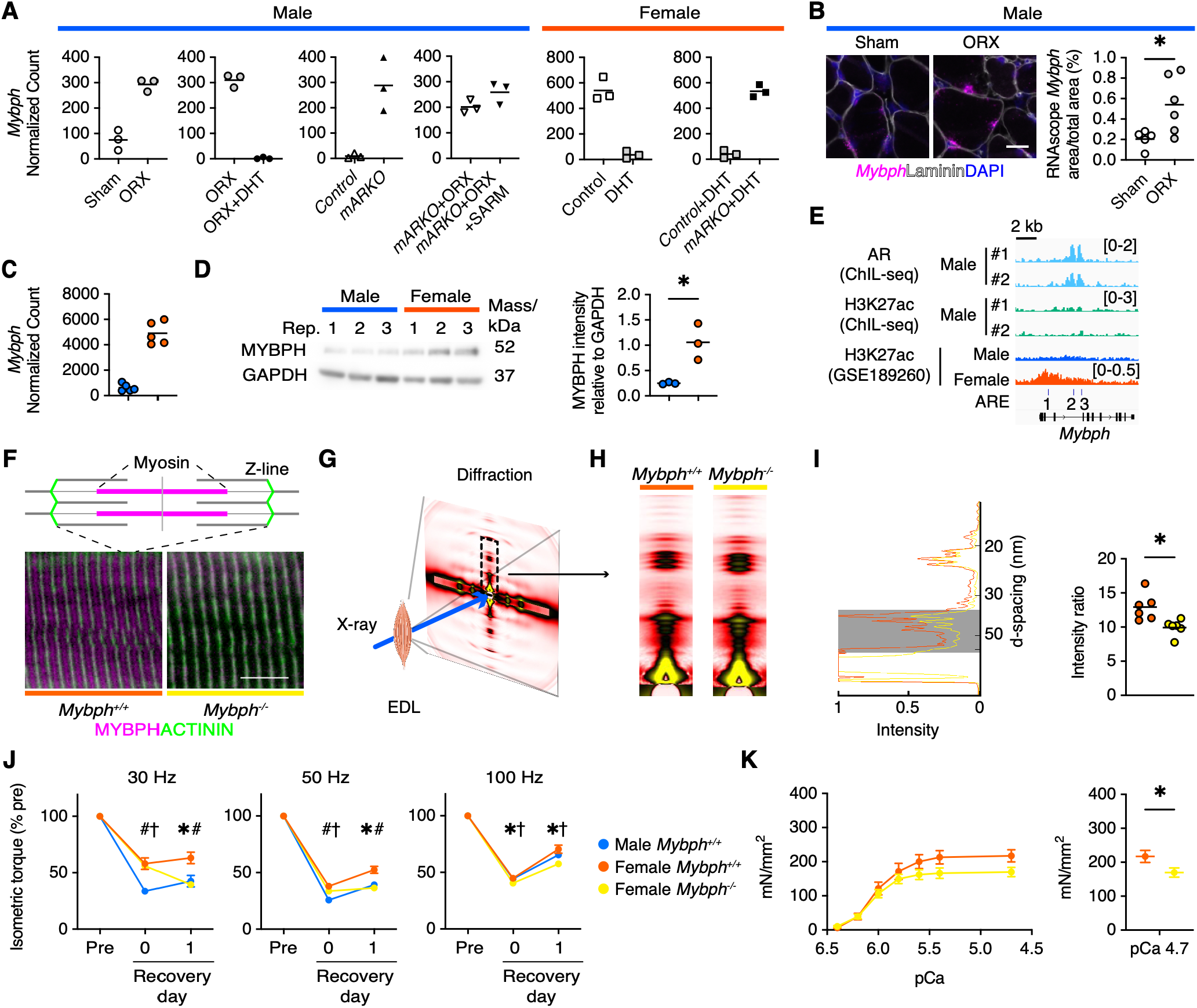
Disrupted myofibrillar alignment and delayed muscle force recovery after injury following loss of the AR target gene *Mybph*. (A) *Mybph* transcripts measured by RNA-seq in androgen-deficient and/or skeletal muscle–specific AR-deficient mice. (B) RNAscope for *Mybph* in skeletal muscle (left) and quantification (right). Scale bar, 25 µm. (C) Comparison of *Mybph* transcripts between male and female skeletal muscle by RNA-seq. (D) Western blot analysis of MYBPH in male and female skeletal muscle (left) and quantification (right). (E) AR binding and histone modifications at the *Mybph* locus. (F) MYBPH distribution in sarcomeres. Scale bar, 10 µm. (G) Schematic of small-angle X-ray diffraction analysis of skeletal muscle. (H) Representative X-ray diffraction patterns. (I) Intensity profiles of meridional reflections in representative X-ray diffraction patterns (left). Area under the curve within the shaded gray region (d = 35 to 70 nm) were calculated and normalized to the M3 intensity (d = 14.3 nm) (right). (J) Measurement of isometric torque before and after ECC-induced injury. (K) Measurement of specific force in skinned fibers one day after ECC injury (left) and measurement at pCa 4.7 (right). ^*^P < 0.05, Welch’s t test. For (J), ^*^P < 0.05 between female *Mybph*^+/+^ and female *Mybph*^-/-^ mice; #P < 0.05 between male *Mybph*^+/+^ and female *Mybph*^+/+^ mice; †P < 0.05 between female *Mybph*^-/-^ and male *Mybph*^+/+^ mice; two-way repeated-measures ANOVA with Tukey’s multiple-comparisons test.

Because snRNA-seq showed that *Mybph* expression was restricted to skeletal muscle fibers (fig. S4A) ^25^, we generated systemic *Mybph*-deficient mice using the CRISPR/Cas9 system to determine the function of MYBPH *in vivo* (fig. S4B). Loss of *Mybph* mRNA and MYBPH protein was confirmed in both male and female knockout mice (fig. S4, C and D). Because *Mybph* was a female-biased gene, we first examined the phenotype of female *Mybph* knockout mice. Subcutaneous white adipose tissue (sWAT) weight was slightly increased in *Mybph* knockout mice (fig. S4E), raising the possibility that systemic metabolism might be altered. However, metabolic assessments, including oxygen consumption (VO_2_), carbon dioxide production (VCO_2_), carbohydrate oxidation rate (CHO), and fat oxidation rate (FAT), showed no significant differences between control and *Mybph* knockout mice (fig. S5, A and B). These findings suggest that the loss of *Mybph* has limited effects on systemic metabolism under basal conditions.

MYBPH localized to the thick filaments of sarcomeres (Fig. 3F), implicating MYBPH in sarcomere organization ^26,27^. To examine sarcomere structure in *Mybph* knockout mice, we next analyzed myofilament organization by small-angle X-ray scattering (Fig. 3G) ^28^. X-ray scattering patterns revealed altered intensities of reflections with d-spacings of 35 to 70 nm, a region containing reflections arising from the axial arrangement of myosin-binding protein C (MyBP-C) ^28^, indicating that loss of Mybph alters sarcomere organization (Fig. 3, H and I, and fig. S6A). By contrast, loss of Mybph did not affect the distribution of myosin heavy chain isoforms (fig. S4F) or the intensities of myosin-based reflections (fig. S6, B to D).

Recovery from contraction-induced muscle damage is faster in female mice than in male mice ^3,4^. To determine whether *Mybph* deficiency affects recovery of contractile function, we measured isometric torque after eccentric contractions (ECC) used to induce muscle damage. Consistent with previous reports, torque recovery was faster in control female mice than in control male mice (Fig. 3J). In female *Mybph* knockout mice, however, recovery was delayed to a level similar to that observed in control male mice (Fig. 3J).

Furthermore, in chemically skinned fiber preparations collected 1 day after eccentric contraction–induced damage, maximal Ca^2+^-activated specific force was reduced in female *Mybph* knockout mice (Fig. 3K and fig. S4G). These findings suggest that *Mybph* deficiency increases susceptibility to eccentric contraction–induced cross-bridge dysfunction. By contrast, male *Mybph*-deficient mice showed no changes in body composition or grip strength (fig. S7A), even after ORX, which increased *Mybph* expression in control male mice (fig. S7B). Together, these findings identify *Mybph* as a sex-biased AR target required for myofibrillar alignment and efficient muscle force recovery in female mice.

### AR regulated *MYBPH* expression in human skeletal muscles

To determine whether AR regulates *MYBPH* in human skeletal muscle, we collected semitendinosus muscle specimens from patients undergoing anterior cruciate ligament (ACL) reconstruction (Fig. 4A). As observed in mice, AR abundance was higher in male than in female human skeletal muscle (Fig. 4, B and C). We then applied AR ChIL-seq to human skeletal muscle specimens (Fig. 4D). AR ChIL-seq identified 6,838 peaks, with the ARE and an AR half-site among the most enriched motifs (Fig. 4E and fig. S8, A and B). Genes associated with ARE-containing AR peaks were enriched for GO Biological Process terms related to “regulation of muscle system processes” (Fig. 4F), consistent with the AR ChIL-seq results from mouse skeletal muscle. AR peaks accompanied by H3K27ac signals were also detected at genes previously identified as androgen- and AR-regulated genes in mouse skeletal muscle (fig. S8C), supporting the applicability of AR ChIL-seq to human skeletal muscle specimens. At the human *MYBPH* locus, we identified an AR peak containing an ARE (Fig. 4G), and this regulatory element was conserved across humans and rodents (Fig. 4H). H3K27ac ChIL-seq further revealed lower signals at the *MYBPH* locus in male muscle than in female muscle (Fig. 4G), consistent with sex-biased repression of *MYBPH* by AR. *MYBPH* transcript abundance was higher in female than in male skeletal muscle (Fig. 4I), in agreement with the findings in mice, and MYBPH protein abundance was likewise higher in female muscle (Fig. 4J). We also confirmed coexpression of NCOR2 and AR in human skeletal muscle fibers (Fig. 4K). Together, these findings support a conserved mechanism by which AR represses *MYBPH* expression in human skeletal muscle.

**Fig 4.**
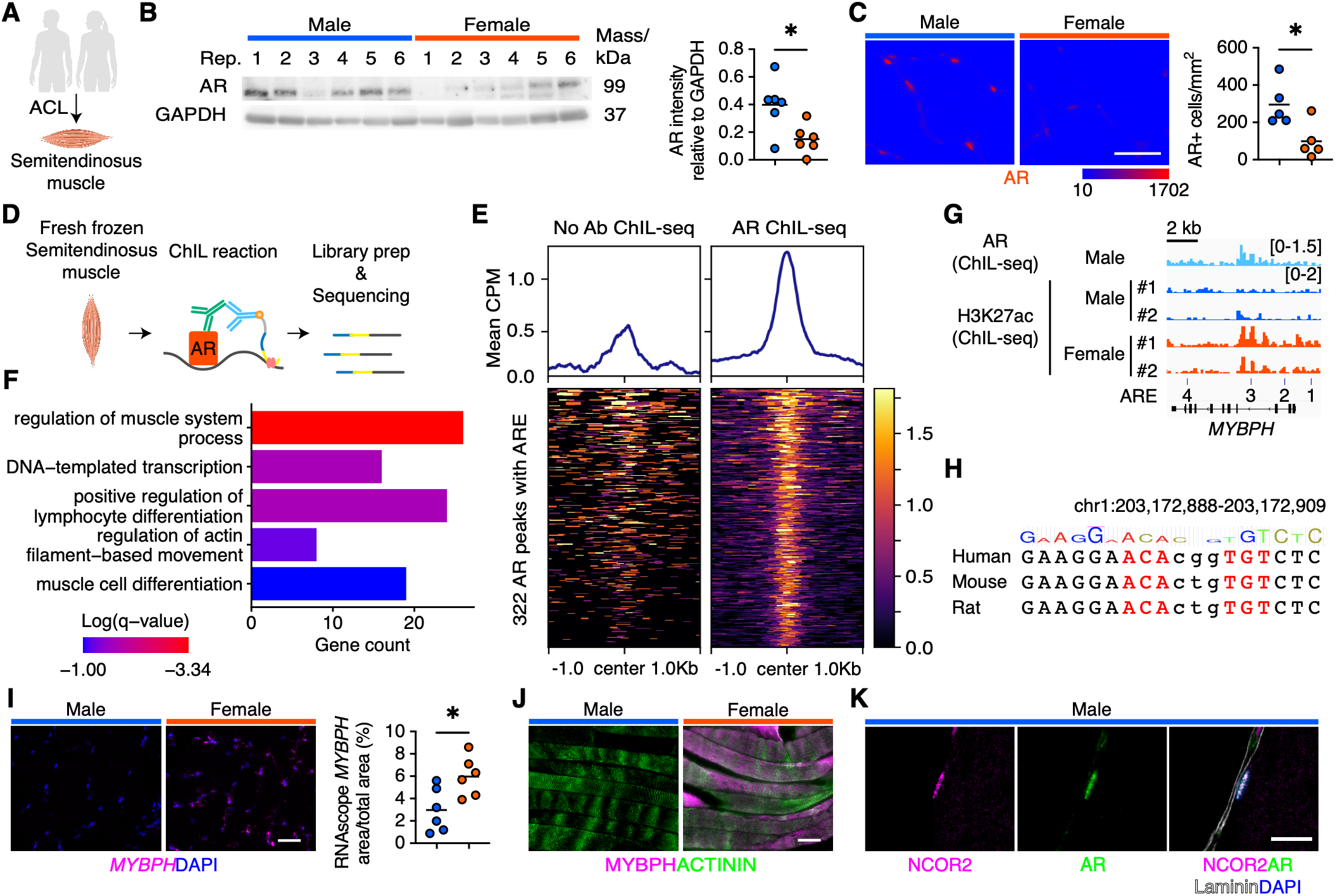
AR-associated *MYBPH* expression in human skeletal muscle. (A) Schematic of human skeletal muscle sampling after ACL reconstruction surgery. ACL, anterior cruciate ligament. (B) Western blot analysis of AR and GAPDH in human skeletal muscle (left) and quantification (right). (C) Immunostaining for AR in human skeletal muscle shown with a BlueToRed LUT (left) and quantification of AR signal (right). Scale bar, 50 µm. (D) Schematic of ChIL-seq analysis in human skeletal muscle. (E) Aggregation plot of AR ChIL-seq signals in human skeletal muscle, centered on AR ChIL-seq peaks containing AREs. CPM, counts per million. (F) Top GO Biological Process terms associated with genes near AR ChIL-seq peaks containing AREs in human skeletal muscle. (G) AR binding and histone modifications detected by ChIL-seq at the *MYBPH* locus in human skeletal muscle. (H) Conserved ARE at the *MYBPH* locus, with the AR ChIL-seq peak shown in (G). (I) RNAscope in situ hybridization for *MYBPH* in human skeletal muscle (left) and quantification (right). Scale bar, 50 µm. (J) Immunostaining for MYBPH and ACTININ in human skeletal muscle. Scale bar, 50 µm. (K) Immunostaining for AR and NCOR2 in human skeletal muscle. Scale bar, 20 µm. ^*^P < 0.05, Welch’s t test.

Previous studies have shown that AR regulates multiple aspects of skeletal muscle physiology through gene programs that vary with cell types and differentiation states. More recently, genome-wide analysis of the AR cistrome in mouse myofibers showed that AR directly activates broad transcriptional programs involved in glycolysis, oxidative metabolism, and muscle contraction ^9^. Our findings not only validate previously defined AR-regulated pathways by recovering *Mylk4* and *Amd1* as androgen-induced ^10,11^, myofiber AR– dependent genes but also extend this framework by identifying *Smtnl2, Slc30a2*, and *Slc38a4* as previously unrecognized targets involved in actin cytoskeletal organization and zinc and amino acid transport. More importantly, by integrating AR occupancy, hormone-responsive H3K27ac, and expression changes following androgen manipulation and myofiber-specific AR deletion, we identified *Mybph* as a high-confidence AR-repressed target with female-biased expression. The conserved AR-binding element and corresponding sex-biased chromatin state and MYBPH expression in mouse and human skeletal muscle support conservation of this regulatory mechanism across species. Together, these findings expand the established role of AR beyond the activation of metabolic and contractile programs and identify AR-dependent transcriptional repression as a potential mechanism contributing to skeletal muscle sexual dimorphism.

Beyond defining AR target genes, understanding how AR produces distinct transcriptional outputs requires identification of its associated coregulators. Most mechanistic insights into AR coregulation have been obtained from prostate-derived cells or artificial expression systems. These studies showed that the corepressors NCOR1 (NCoR) and NCOR2 (SMRT) interact with AR and suppress its transcriptional activity, whereas LSD1 directly associates with AR and can function as either a coactivator or a corepressor, depending on the target gene and chromatin context ^21,22^. Our AirID-AR knock-in model extends these observations to endogenous AR in skeletal muscle *in vivo* under physiological condition, identifying established AR-associated proteins and NCOR2-linked histone deacetylase machinery that may mediate AR-dependent repression. Biotinylation of NCOR2 within its nuclear receptor–interaction region, together with the coexpression of AR and NCOR2 in mouse and human myofibers, further supports a physiologically relevant association. By contrast, LSD1 was not detected, suggesting that the AR-associated proteome in normal skeletal muscle differs substantially from those characterized in prostate tissue and cancer-derived cells.

MYBPH has long been recognized as a myosin-associated component of the sarcomeric C-zone ^26,27^, but its physiological function in adult mammalian skeletal muscle has remained largely unknown. Recent work in zebrafish show that Mybph competes with fast skeletal Mybpc for C-zone occupancy and restrains actin sliding, although *Mybph*-deficient larvae show no overt basal muscle phenotype, possibly because of compensation by Mybpc ^29^. Similarly, in our study, *Mybph*-deficient mice were largely normal under basal conditions, whereas small-angle X-ray scattering revealed altered organization of the Mybp-containing region, and eccentric contractions unmasked reduced maximal Ca^2+^-activated specific force and delayed force recovery in females. These findings suggest that MYBPH is largely dispensable for basal muscle function but becomes important for maintaining sarcomeric organization and cross-bridge function under mechanical stress.

Previous studies showed that female mice recover force more rapidly after eccentric contractions and recover muscle size and function more rapidly after toxin-induced injury ^3–5^. This advantage has often been attributed to the proposed membrane-stabilizing and anti-inflammatory effects of estrogen ^30,31^, although evidence remains mixed, as estradiol replacement did not protect ovariectomized mice from acute eccentric contraction–induced injury ^32^. Instead, we identify a myofiber-intrinsic androgen–AR–MYBPH axis through which androgen signaling suppresses a sarcomeric factor that contributes to the recovery of cross-bridge function after muscle injuruy. Conservation of AR occupancy, sex-biased chromatin acetylation, and MYBPH expression in human muscle suggests that this mechanism may contribute to human sexual dimorphism, although its role in human muscle injury remains to be established. Together, these results identify MYBPH as a sex-biased AR target that may contribute to sex differences in postinjury muscle force recovery and suggest that androgen-dependent repression of MYBPH acts alongside estrogen-mediated protection in shaping this dimorphism.

Together, our study defines an *in vivo* AR regulatory network in skeletal muscle that integrates genomic targets, associated coregulators, and a sex-biased effector of postinjury contractile recovery. More broadly, the integration of ChIL-seq with AirID provides a generalizable strategy for defining tissue-specific hormone receptor networks and identifying molecular effectors of sexual dimorphism in other tissues. By linking AR-dependent MYBPH repression to sex-biased muscle function in mice and showing conservation of this regulatory axis in human muscle, these findings provide a mechanistic framework for understanding how androgen signaling contributes to skeletal muscle sexual dimorphism.

## Acknowledgments

We are grateful to Y. Sato and M. Chosei for technical support. We thank K. Ojima of the Division of Animal Products Research, NARO Institute of Livestock and Grassland Science (NILGS), for advice on immunostaining. The synchrotron radiation experiments were performed at the BL05XU and BL40XU of SPring-8 with the approval of the Japan Synchrotron Radiation Research Institute (JASRI) (Proposal No. 2025A1114, 2025B1294, and 2026A1418). We also thank S. Kamei at Minami Matsuyama Hospital and Y. Taguchi at Okujima Hospital for assistance with human skeletal muscle sampling. We thank N. Tokunaga for library preparation and the members of the Division of Medical Research Support, Advanced Research Support Center, Ehime University, for technical assistance. This work was also supported in part by the MEXT Promotion of Development of a Joint Usage/Research System Project: Pan-Omics DDRIC, the Medical Research Center Initiative (MRCI) for High Depth Omics, CURE (JPMXP1323015486) for MIB and RIIT, and AMRC at Kyushu University. ChatGPT-5.5 was used to assist with language editing and improving the clarity of the manuscript.

## Author contributions

Conceptualization: HS, YI

Methodology: HS, YY, KT, AT, HI, YH, SO, TK, KW, KH, MT, PM, ST, HK, TS, TY, YK, AH, YO, YI

Investigation: HS, YY, KT, AT, HI, HK, TY, AH

Visualization: HS, YY, KT, AT, HI, HK, TY, AH

Funding acquisition: HS, YY, TY, YK, YI

Project administration: HS, YI

Supervision: HS, YI

Writing – original draft: HS, YY, YI

Writing – review & editing: HS, YY, AT, HI, HK, TY, YI

## Data, code, and materials availability

RNA-seq and ChIL-seq data have been deposited and are publicly available in the Gene Expression Omnibus as accession number GSE341550 and GSE341548.

## Supplementary Materials

Materials and Methods

Figs. S1 to S8

Table S1

Data S1 to S4

