## Supplemental Figures for "Androgen receptor determines skeletal muscle sexual dimorphism"

**The PDF file includes:**

Materials and Methods
Figs. S1 to S8
Table S1
Captions for Data S1 to S4
References

**Other Supplementary Materials for this manuscript include the following:**

Data S1 to S4

### Materials and Methods

Mice

C57BL/6J male mice were purchased from Jackson Laboratories (Cat# 000664, RRID:IMSR_JAX:000664). The mice were maintained in a controlled environment within a specific pathogen-free facility, with climate-controlled conditions and a 12-hour light-dark cycle. Mice had unrestricted access to water and were fed a standard diet. For RNA-seq, mice were randomized based on body weight and subjected to orchiectomy (ORX) under anaesthesia. On the same day, dihydrotestosterone (DHT; stanolone, 2.5 mg/mouse; Tokyo Chemical Industry, Cat# A0462), prepared in absolute ethanol and diluted in PBS containing 0.3% (w/v) hydroxypropyl cellulose (Wako, Cat# 085-07932), was injected subcutaneously. The care and usage of all animals in this study were conducted in strict adherence to the guidelines and protocols approved by the Animal Experiment Committee of Ehime University, Japan.

Generation of genetically modified mice using CRISPR/Cas9

AirID-AR knock-in mice and systemic Mybph knockout mice were generated using the CRISPR/Cas9 system. Single-guide RNAs (sgRNAs) targeting first ATG of the mouse *Ar* (Gene ID: 11835) locus and exon 1 of the mouse *Mybph* (Gene ID: 53311) locus were designed using Benchling. sgRNA were synthesized using Precision gRNA Synthesis Kit (Invitrogen, Cat# A29377) according to the manufacturer's instructions. Recombinant Cas9 protein (TrueCut™ Cas9 Protein v2, Invitrogen, Cat# A36499) was complexed with sgRNA to form Cas9–gRNA ribonucleoprotein (RNP) complexes. The RNP complexes were introduced by electroporation into fertilized one-cell-stage embryos obtained from C57BL/6J mice purchased from Jackson Laboratory Japan. Electroporation was performed under the following conditions: 25 V, 3 ms ON, 97 ms OFF, 5 pulses [1–3].

For the generation of AirID-AR mice, following electroporation, the embryos were incubated with a custom single-stranded adeno-associated virus serotype 6 (ssAAV6) donor vector carrying the desired cassette flanked by homology arms. The ssAAV6 donor vector was produced by VectorBuilder Japan and was used as the donor template for homology-directed repair [4]. For the generation of systemic *Mybph* knockout mice, the Cas9–sgRNA RNP complexes were introduced into fertilized one-cell-stage embryos by electroporation without a donor template. Treated embryos were cultured to the two-cell stage and transferred into the oviducts of pseudopregnant recipient female Jcl:ICR mice purchased from CLEA Japan.

Tissue chromatin integration labelling (ChIL) experiments

For ChIL-seq, C57BL/6J male mice were treated with DHT 2h before sampling to enhance AR expression in skeletal muscles [5]. ChIL-seq analysis was performed as previously described [6,7], using anti-AR (Abcam, Cat# ab108341, RRID:AB_10865716; 1/100) and anti-H3K27ac (MAB, Cat# MABI0309, RRID:AB_11126964) antibodies. Sequencing of the libraries was performed using the Hiseq1500 and NovaSeq 6000 platforms (Illumina).

To align the reads, the GRCm38 reference genomes were employed, utilizing Bowtie2 with the default parameters [8]. Removal of duplicated reads was carried out using Samtools (rmdup) [9]. Subsequent analysis focused solely on the uniquely mapped reads. To identify AR peaks across samples, peak calling was performed with MACS2 on BAM files merged from biological replicates ($n = 4$). The MACS2 algorithm with the default parameters [10] was employed for peak calling, using ChIL probe-only sections (for AR) or non-treated muscle section (for H3K27ac) as the control. To ensure high-quality peaks, any peaks with $p < 0.05$ (called for AR with ChIL probe-only control) or a false discovery rate (FDR) exceeding 0.05 (H3K27ac) were excluded from subsequent analysis.

Heatmaps of AR and H3K27ac ChIL-seq signals were generated using deepTools plotHeatmap based on CPM-normalized bigWig files pooled across replicates [11]. The distribution of mapped reads across genomic features was determined using HOMER (annotatePeaks.pl) [12]. Motif analysis was performed using HOMER with the following command: findMotifsGenome.pl -size 200 -mask. Pearson correlation coefficients were calculated using deepTools by summarizing BAM files with multiBamSummary and then performing correlation analysis with plotCorrelation -c pearson --skipZeros --removeOutliers. Publicly available ChIP-seq data were obtained from the NCBI Gene Expression Omnibus, including GSE47192, which contains AR ChIP-seq data from the prostate, kidney, and epididymis.

Enrichment of biotinylated peptides

AirID-AR mice were administered 2 mM biotin (Nacalai Tesque, Cat# 04822-91) orally for 3 days before sample collection. DHT was injected subcutaneously on the first day of biotin administration. Tibialis anterior (TA) muscles were collected, and biotinylated peptides were enriched as described previously [13]. Briefly, muscle lysates were prepared in Gdm-TCEP buffer consisting of 6 M guanidine hydrochloride, 100 mM HEPES–NaOH (pH 7.5), 10 mM tris(2-carboxyethyl)phosphine, and 40 mM chloroacetamide. The lysates were heated, sonicated, and centrifuged at 20,000 × g for 15 min at 4 °C. The supernatants were collected, and the proteins were purified by methanol–chloroform precipitation and resuspended in 150 µL of phase-transfer surfactant (PTS) buffer consisting of 12 mM sodium deoxycholate, 12 mM sodium lauroyl sarcosinate, and 100 mM Tris–HCl (pH 8.0). After further sonication and heating, the protein solutions were diluted fivefold with 100 mM Tris–HCl (pH 8.0) and digested overnight at 37 °C with MS-grade trypsin (Thermo Fisher Scientific, Cat# 90057). The resulting peptide solutions were diluted twofold with TBS consisting of 50 mM Tris–HCl (pH 7.5) and 150 mM NaCl. Biotinylated peptides were enriched by incubating the samples with 15 µL of MagCapture HP Tamavidin 2-REV magnetic bead slurry (FUJIFILM Wako, Cat# 133-18611) for 3 h at 4 °C. After five washes with TBS, the bound biotinylated peptides were eluted twice with 100 µL of 1 mM biotin in TBS for 15 min at 37 °C. The eluates were combined, desalted using a GL-Tip SDB cartridge (GL Sciences, Cat# 7820-11200), dried in a SpeedVac concentrator, and reconstituted in 0.1% trifluoroacetic acid containing 3% acetonitrile.

Data-dependent LC-MS/MS analysis

Biotinylated peptides were analyzed using an EASY-nLC 1200 UHPLC system coupled to an Orbitrap Fusion mass spectrometer through a nanoelectrospray ion source (Thermo Fisher Scientific). Peptides were separated on a C18 reversed-phase analytical column (150 mm × 75 µm inner diameter; Nikkyo Technos) using a linear gradient from 4% to 32% acetonitrile over 60 min, followed by an increase to 80% acetonitrile over 10 min and a further 10-min hold at 80% acetonitrile. The mass spectrometer was operated in data-dependent acquisition mode with a maximum cycle time of 3 s. Full MS spectra were acquired in the Orbitrap at a resolution of 120,000, with an automatic gain control target of $4 \times 10^5$ and an m/z range of 375–1,500. HCD MS/MS spectra were acquired in the linear ion trap with an automatic gain control target of $1 \times 10^4$, an isolation window of 1.6 m/z, a maximum injection time of 200 ms, and a normalized collision energy of 30. Dynamic exclusion was set to 10 s. Raw data were searched against the Swiss-Prot database restricted to Mus musculus using Proteome Discoverer version 2.5 (Thermo Fisher Scientific) with the Sequest HT search engine. Trypsin was specified as the proteolytic enzyme, and up to two missed cleavages were allowed. The precursor and fragment mass tolerances were set to 10 ppm and 0.6 Da, respectively. Carbamidomethylation of cysteine was specified as a fixed modification, whereas acetylation of the protein N-terminus, oxidation of methionine, and biotinylation of lysine were specified as variable modifications. Peptide identifications were filtered at a false discovery rate of 1% using the Percolator node. Label-free quantification was performed on the basis of precursor ion intensities using the Precursor Ions Quantifier node. Abundance values were normalized such that the total peptide abundance was equal across samples. Statistical analyses were performed using Microsoft Excel version 16.78.3. P values were calculated using Student's t-test and adjusted for multiple comparisons using the Benjamini–Hochberg procedure. The adjusted P values are provided in Tables S2 and S4. Gene Ontology enrichment analysis was performed using Metascape [14], and the results were visualized using R.

Immunofluorescence staining and microscopy

For immunofluorescence analysis, muscles were rapidly frozen in isopentane prechilled with liquid nitrogen. Cryosections (10 μm) were then prepared for staining. Sections were fixed with 4% paraformaldehyde in PBS for 5 min, blocked with 5% goat serum (Gibco, Cat# 16210-064) in PBS or Blocking One Histo (Nacalai Tesque, Cat# 06349-64) for 60 min at room temperature, and subsequently incubated with primary antibodies overnight at 4°C. For myosin heavy chain (Myh) staining, sections were first air-dried and then blocked with M.O.M. reagent (Vector Laboratories, Cat# MKB-2213-1) at room temperature for 60 min. Primary antibodies against Myh were diluted in 1% BSA/PBS and applied for 45 min at 37°C. The antibodies used for immunofluorescence staining were as follows: anti-AR (Abcam, Cat# ab108341, RRID:AB_10865716; 1:100), anti-MYBPH (Sigma-Aldrich, Cat# HPA061383, RRID:AB_2684497; 1/150), anti-Actinin (Sigma-Aldrich, Cat# A7811, RRID:AB_476766; 1/400), anti-Laminin α2 (Santa Cruz Biotechnology, Cat# sc-59854, RRID:AB_784266; 1:400), anti-Laminin (Sigma-Aldrich, Cat# L9393, RRID:AB_477163; 1:500), anti-Myh7 (DSHB, Cat# BA-D5, RRID:AB_2235587, supernatant; 1:100), anti-Myh2 (DSHB, Cat# SC-71, RRID:AB_2147165, supernatant; 1:100), anti-Myh4 (DSHB, Cat# BF-F3, RRID:AB_2266724, concentrated purified; 1:100), Alexa Fluor 568 goat anti-mouse IgG1 (Thermo Fisher Scientific, Cat# A-21124, RRID:AB_2535766; 1:1000), Alexa Fluor 488 goat anti-rabbit IgG (Thermo Fisher Scientific, Cat# A-11008, RRID:AB_143165; 1:1000), Alexa Fluor 488 goat anti-mouse IgM (Thermo Fisher Scientific, Cat# A-21042, RRID:AB_143165; 1:1000), and Alexa Fluor 647 goat anti-rabbit IgG (Thermo Fisher Scientific, Cat# A27040, RRID:AB_143165; 1:1000). Nuclei were counterstained with DAPI.

To quantify myofiber number, cross-sectional area (CSA), and minimal Feret diameter, images of entire muscle cross-sections were analyzed using Fiji (v2.3.0) (https://imagej.net/Fiji). Myofiber areas were identified based on laminin-positive signals, and regions of interest (ROIs) were defined within the laminin-positive areas. The frequency of each muscle fiber type within the ROI was determined using the Trainable Weka Segmentation plugin in Fiji after manual training. Small ($<150\ \mu m^2$) and large ($>10{,}000\ \mu m^2$) fibers were excluded from the analysis. The numbers of AR-positive cells were manually counted using Fiji.

RNA-seq

TA muscles were collected from sham-operated, ORX, and ORX+DHT-treated mice. Total RNA was isolated from the muscles by Isogen (Nippon Gene, Cat# 311-07361) according to the manufacturer's instructions. Libraries were prepared using the NEBNext Ultra II Directional RNA Library Prep Kit for Illumina (New England Biolabs, Cat# E7760), together with the NEBNext Poly(A) mRNA Magnetic Isolation Module (New England Biolabs, Cat# E7490) and NEBNext Multiplex Oligos for Illumina (New England Biolabs,

Cat# E7335). Library quality was assessed using a Bioanalyzer with the DNA 1000 Kit (Agilent, Cat# 5067-1504), and library concentration was determined using the KAPA Library Quantification Kit (Roche, Cat# KK4824). Libraries were sequenced on a MiSeq instrument using the MiSeq Reagent Kit v3 (150 cycles) to generate 75-bp paired-end reads.

Adapter sequences and low-quality bases were removed using Trim Galore (v0.6.7, www.trimgalore.com/), and reads shorter than 20 bp were discarded. The processed reads were aligned to the GRCm38 reference genome using HISAT2 (v2.2.1) [15]. Gene-level read counts were generated using featureCounts (v2.0.1) [16]. Differential expression analysis was performed using DESeq2 (v1.36.0) [17], and genes with |log2FC| > 1 and padj < 0.1 were considered differentially expressed. Publicly available RNA-seq data were obtained from the NCBI Gene Expression Omnibus, including GSE153147 for male myofiber-specific AR knockout mice and GSE152756 for female myofiber-specific AR knockout mice, and the Sequence Read Archive database under the BioProject accession number PRJNA1172958 for sex dimorphism of gene expression in mice.

<u>RNAscope in situ hybridization and quantification on muscle sections</u>

RNAscope in situ hybridization was performed on mouse and human skeletal muscle sections as described previously [18,19]. Cryosections (10 µm) were baked at 55°C, washed in PBS, and treated with hydrogen peroxide at room temperature for 10 min. After target retrieval at 95°C with Target Retrieval Reagents (ACD, Cat# 322000), sections were dehydrated in 100% ethanol, dried, and outlined with a hydrophobic barrier (Vector Laboratories, Cat# H-4000). Sections were then treated with Protease Plus for 15 min at 37°C, followed by hybridization with RNAscope probes against *Mybph* for 2 h at 37°C. After washing, signal amplification was performed according to the RNAscope Multiplex Fluorescent Reagent Kit v2 workflow, followed by HRP-based detection with Opal 570 fluorophores (Akoya Biosciences, Cat# OP-001003; 1/1500). For sections combined with immunofluorescence staining, slides were subsequently blocked in PBS containing 5% goat serum, incubated with primary antibodies against Laminin overnight at 4°C, and then with appropriate secondary antibodies and DAPI.

Entire cross-sectional fluorescence images stained for *Mybph* and laminin were acquired using an Axio Observer 7 microscope (Zeiss). The *Mybph*-positive area was binarized and quantified using Fiji. The total cross-sectional area was defined on the basis of laminin staining.

<u>Western blotting</u>

TA muscles were homogenized in RIPA buffer (Fujifilm, Cat# 182-02451) supplemented with protease inhibitors (Nacalai Tesque, Cat# 25955-11). Tissue extracts were separated by SDS-PAGE and transferred onto polyvinylidene fluoride membranes. Membranes were blocked with 5% skim milk in TBS containing Tween-20 (TBST), followed by incubation overnight at 4°C with anti-AR, anti-MYBPH (Sigma-Aldrich, Cat# HPA061383, RRID:AB_2684497; 1/150) or anti-GAPDH (Cell Signaling Technology, Cat# 5174, RRID:AB_10622025; 1:1000) antibodies. Membranes were then incubated for 60 min at room temperature with HRP-conjugated anti-mouse (Agilent, Cat# P0260, RRID:AB_2636929; 1:5000) or anti-rabbit immunoglobulin secondary antibodies (Agilent, Cat# P0448, RRID:AB_2617138; 1:5000). Signals were detected using ECL Prime Western Blotting Detection Reagents (Amersham, Cat# RPN2232) and imaged with an ImageQuant LAS 4000 system (GE Healthcare). Band intensities were quantified using Fiji.

<u>Muscle torque measurement, eccentric contractions, and skinned fiber analysis</u>

Mice anesthetized with 2% isoflurane were positioned on a platform so that one hindpaw was aligned at a 90° angle to the tibia on a sensor footplate. The plantar flexor muscles were stimulated supramaximally (45 V) through a pair of surface electrodes under isoflurane anesthesia in female *Mybph*$^{+/+}$ and *Mybph*$^{-/-}$, and male *Mybph*$^{+/+}$ mice. Eccentric contractions (ECCs) were induced 100 times at 4-s intervals by forced dorsiflexion from 0° to 40° at an angular velocity of 150°/s in combination with electrical stimulation (0.5-ms monophasic rectangular pulses at 50 Hz), as described previously [20]. Isometric torque was measured at stimulation frequencies of 30, 50, and 100 Hz before, immediately after, and 1 day after the ECC protocol.

Chemically skinned muscle fibers were prepared, and $Ca^{2+}$-activated force was measured as described previously [21]. The gastrocnemius muscle was pinned out at resting length under paraffin oil kept at 4˚C. Single muscle fibers were dissected under a stereomicroscope. Four to six skinned fibers were obtained from each muscle. A segment of each skinned fiber was connected to a force transducer (Muscle Tester, World Precision Instruments) and then incubated in a *N*-2-hydroxyethylpiperazine-*N*'-2-ethanesulfonic acid (HEPES) buffered solution (see below) containing 1% (vol/vol) Triton X-100 for 10 min in order to remove membranous structures. Fiber length was adjusted to the optimal length (2.5 μm) by laser diffraction as described previously [22] and the contractile properties were measured at room temperature (24˚C).

All solutions were prepared as described in detail elsewhere [23]. They contained (in mM) 36 $Na^+$, 126 $K^+$, 90 HEPES, 8 ATP and 10 creatine phosphate, and had a pH of 7.09–7.11 and a free $Mg^{2+}$ concentration of 1.0 mM. The maximum $Ca^{2+}$ solution contained 49.5 mM Ca-EGTA and 0.5 mM free EGTA, whereas the relaxation solution contained 50 mM free EGTA. Various pCa (-log10 of the free $Ca^{2+}$ concentration) solutions (pCa 6.4, 6.2, 6.0, 5.8, 5.6, 5.4, and 4.7) were prepared by mixing the maximum $Ca^{2+}$ solution and the relaxation solution in appropriate proportions [24]. The contractile apparatus was directly activated by exposing the skinned fibers to the various pCa solutions, and force was measured. The cross-sectional area of each fiber was calculated from its diameters. $Ca^{2+}$-activated force normalized to the fiber cross-sectional area is expressed as $mN/mm^2$.

<u>Small-angle X-ray diffraction</u>

All small-angle X-ray diffraction experiments were performed at beamlines BL05XU and BL40XU of SPring-8. *Mybph*$^{+/+}$ and *Mybph*$^{-/-}$ mice were euthanized by cervical dislocation under deep anesthesia. The extensor digitorum longus (EDL) muscles were then excised and immediately mounted in a measurement chamber filled with physiological saline. The chamber was positioned on a motorized XZ sample stage at the beamline, and X-ray diffraction patterns were recorded from the muscles under resting conditions. The incident X-ray energy was 12.4 keV, corresponding to a wavelength of 0.1 nm. A softly focused beam with approximate horizontal and vertical dimensions of 0.15 and 0.10 mm, respectively, was used at the sample position. The sample-to-detector distance was 2m for determining the intensity ratio of the sixth actin layer-line reflection to the 14.3-nm myosin meridional reflection (M3), and 4 m for more precise mesurements of the central region of the diffraction pattern. Diffraction images were acquired using a 4-inch image intensifier (Model V7739PMOD, Hamamatsu Photonics, Japan) coupled to an ORCA-Flash4.2 camera (Hamamatsu Photonics). A 4-mm-diameter flying beamstop was used to block the direct beam. The X-ray diffraction patterns recorded under the same experimental conditions were summed, the four quadrants were averaged, and the background scattering was subtracted as described [25].

<u>Reanalysis of single-cell RNA-seq</u>

Limb muscle scRNA-seq data were obtained from Tabula Muris (droplet, Limb_Muscle-10X_P7_14) and loaded into Seurat. Data were analyzed using the standard Seurat preprocessing workflow. Cells with fewer than 200 or more than 2,500 detected features were excluded, as were cells with >5% mitochondrial counts. Data were normalized using normalization.method = "LogNormalize" and scale.factor = 10000. Clustering was performed with FindNeighbors(dims = 1:10) and FindClusters(resolution = 0.1). To identify *Mybph*+ clusters, dot plots were generated using the following marker genes: Pecam1, Pdgfra, Cd79a, Cd3g, Myod1, Scx, Lyz1, Acta2, Acta1, and Mybph.

Metabolic assessments

Metabolic assessments were performed as previously described [26]. Oxygen consumption (VO2) and carbon dioxide production (VCO2) were measured using a metabolic gas analysis system (Arco Systems, Cat# ARCO-2000). Mice were housed individually in metabolic chambers with free access to food and water. After 3 days of acclimation, measurements were collected from each mouse. Data were recorded every 10 min, and hourly averages were used for analysis. Carbohydrate (CHO) and fat (FAT) oxidation rates were calculated using Frayn's equation, and energy expenditure was calculated using Lusk's equation. Data were normalized to body weight. Locomotor activity was monitored using the Actracer 20000 SN (Arco Systems).

Samling of human muscle samples

All experiments involving human samples were approved by the Institutional Review Board of Ehime University (approval no. IRB2502004), and written informed consent was obtained after the nature and possible consequences of the study had been fully explained. Muscle samples were collected from the semitendinosus muscle during anterior cruciate ligament (ACL) reconstruction surgery. The excised samples were immediately frozen in isopentane prechilled with liquid nitrogen for immunostaining or snap-frozen in liquid nitrogen for western blot analysis.

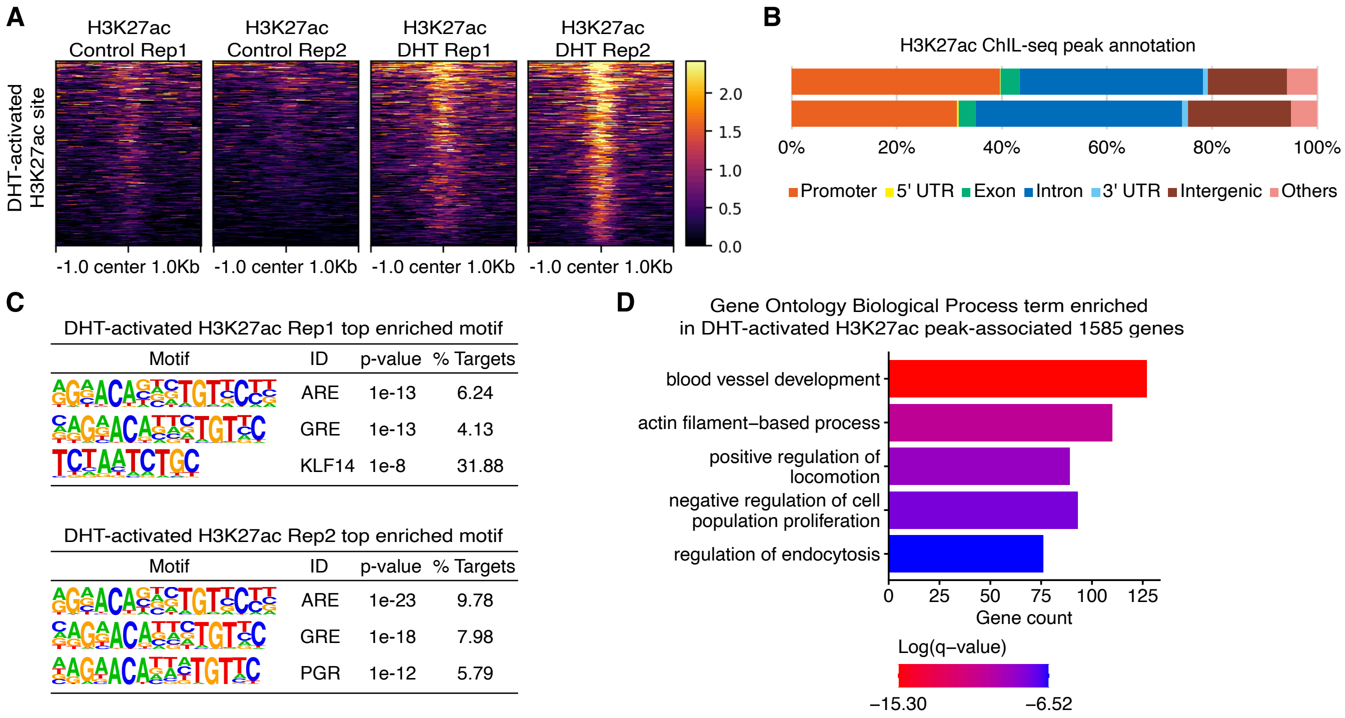


**Fig. S1. Analysis of skeletal muscle H3K27ac ChIL-seq data set.**
(A) Heat map of H3K27ac ChIL-seq signal, shown as CPM, across ±1-kb regions centered on DHT-induced H3K27ac peaks. (B) Genomic distribution of DHT-induced H3K27ac peaks. (C) Enriched motifs at DHT-activated H3K27ac sites. (D) Gene Ontology enrichment analysis of genes associated with DHT-activated H3K27ac sites.

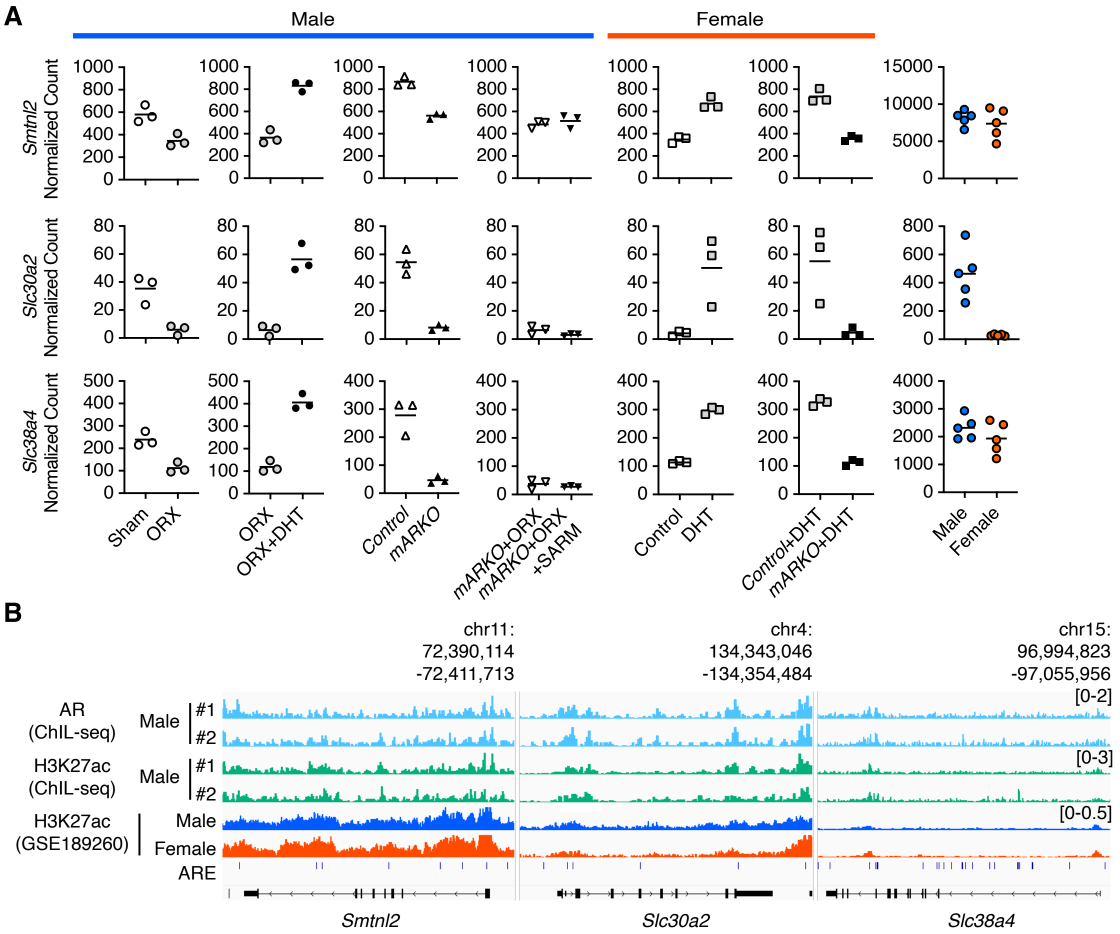


**Fig. S2. Integrated analysis of ChIL-seq and RNA-seq.**
(A) Expression patterns of genes regulated by androgens and AR in skeletal muscle fibers.
(B) AR bindings and histone modifications at androgen- and AR-regulated genes in skeletal muscle fibers.

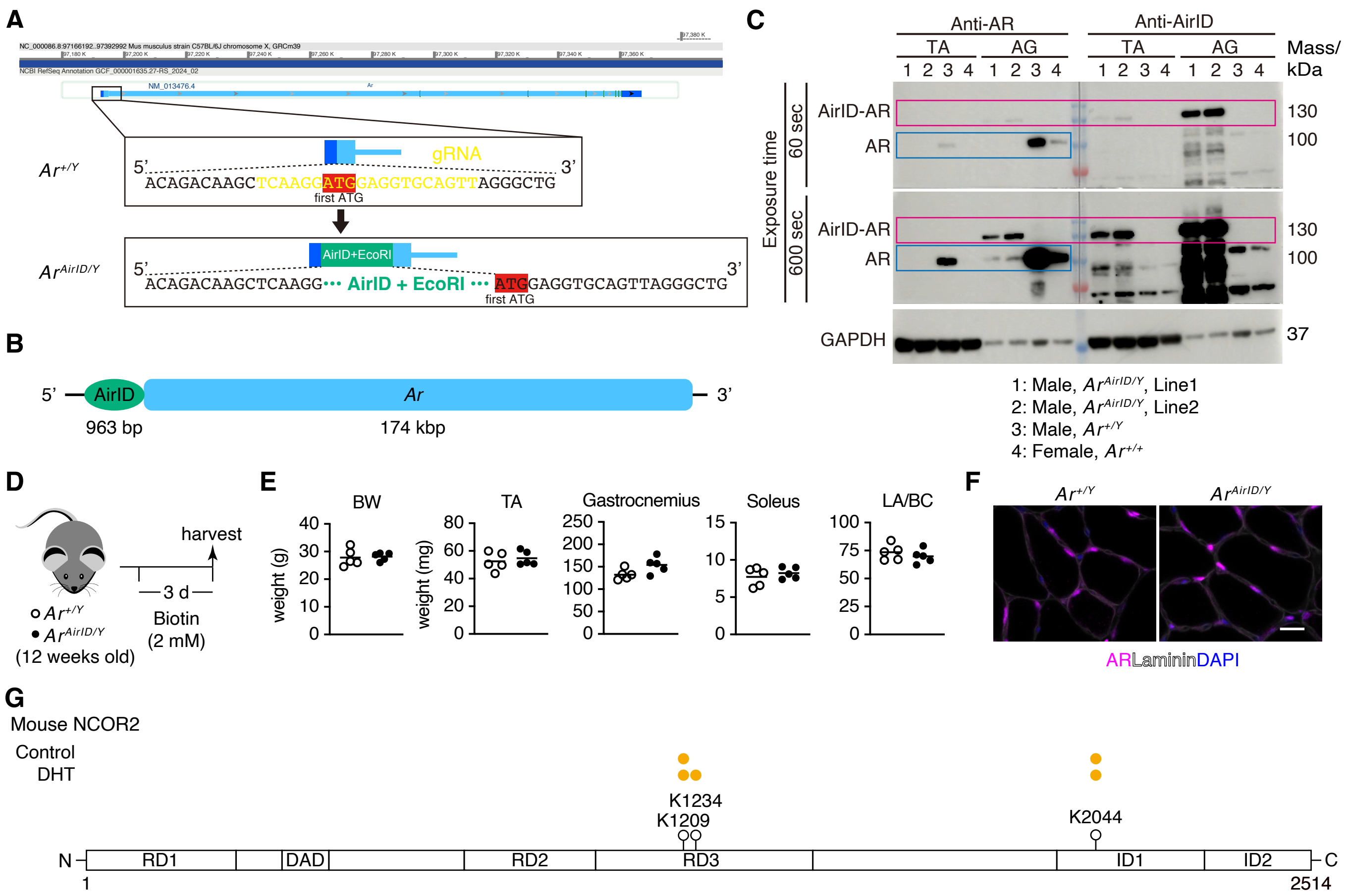


**Fig. S3. Generation of AirID-AR mice.**
(A) Generation of AirID-AR mice, in which AirID was inserted at the first ATG (red) of the endogenous *Ar* locus. (B) Schematic of the *Ar* transcript fused to AirID. (C) Western blot analysis of AirID-AR protein in skeletal muscle and adrenal gland from control and AirID-AR mice using anti-AR and anti-AirID antibodies. (D) Experimental scheme for biotin administration to AirID-AR mice. (E) Quantification of body weight and muscle mass. BW, body weight; TA, tibialis anterior; LA, levator ani; BC, bulbocavernosus; n = 5 mice per genotype. (F) Immunofluorescence for AR and laminin, with DAPI counterstaining, in TA muscles from control and AirID-AR mice. Scale bar, 25 µm. (G) Biotinylated sites detected in NCOR2 from AirID-AR mice with or without DHT treatment. Orange dots denote biotinylated lysine (K) residues.

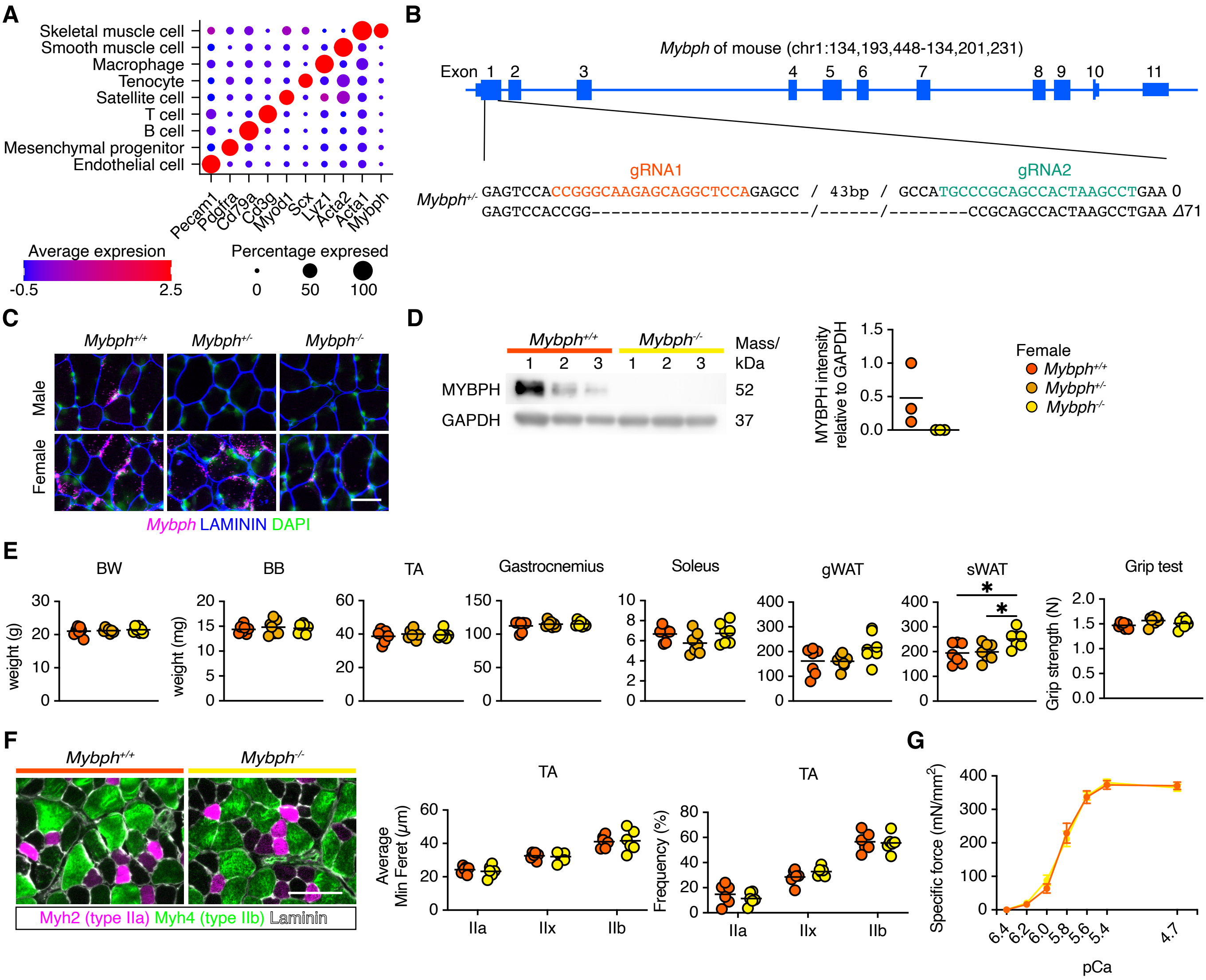


**Fig. S4. Generation and phenotypic analysis of *Mybph* knockout mice.**
(A) *Mybph* expression in skeletal muscle, analyzed using single-cell RNA sequencing data from the Tabula Muris data set. (B) Generation of *Mybph* knockout mice by CRISPR-Cas9. (C) RNAscope in situ hybridization for *Mybph* transcripts in TA muscles from control, heterozygous, and homozygous *Mybph* knockout mice. Scale bar, 100 μm. (D) Immunoblot analysis of MYBPH in TA muscles from female control and *Mybph* knockout mice (left) and quantification (right; n = 3 mice per genotype). (E) Quantification of body weight (BW), muscle mass, gonadal white adipose tissue (gWAT) and subcutaneous white adipose tissue (sWAT) mass, and grip strength in 12-week-old female control, *Mybph* heterozygous, and *Mybph* knockout mice (n = 6 mice per genotype). BB, biceps brachii; TA, tibialis anterior. (F) Immunofluorescence for fast-twitch myosin heavy chain isoforms MYH2 (type IIa) and MYH4 (type IIb), together with laminin, in TA muscles from female control and *Mybph* knockout mice (left), and quantification of mean minimum Feret diameter and fiber-type frequency (right; n = 6 mice per genotype). Scale bar, 100 μm. (G) Quantification of specific force in skinned fibers isolated from intact gastrocnemius muscles of female control and *Mybph* knockout mice (n = 5 mice per genotype).

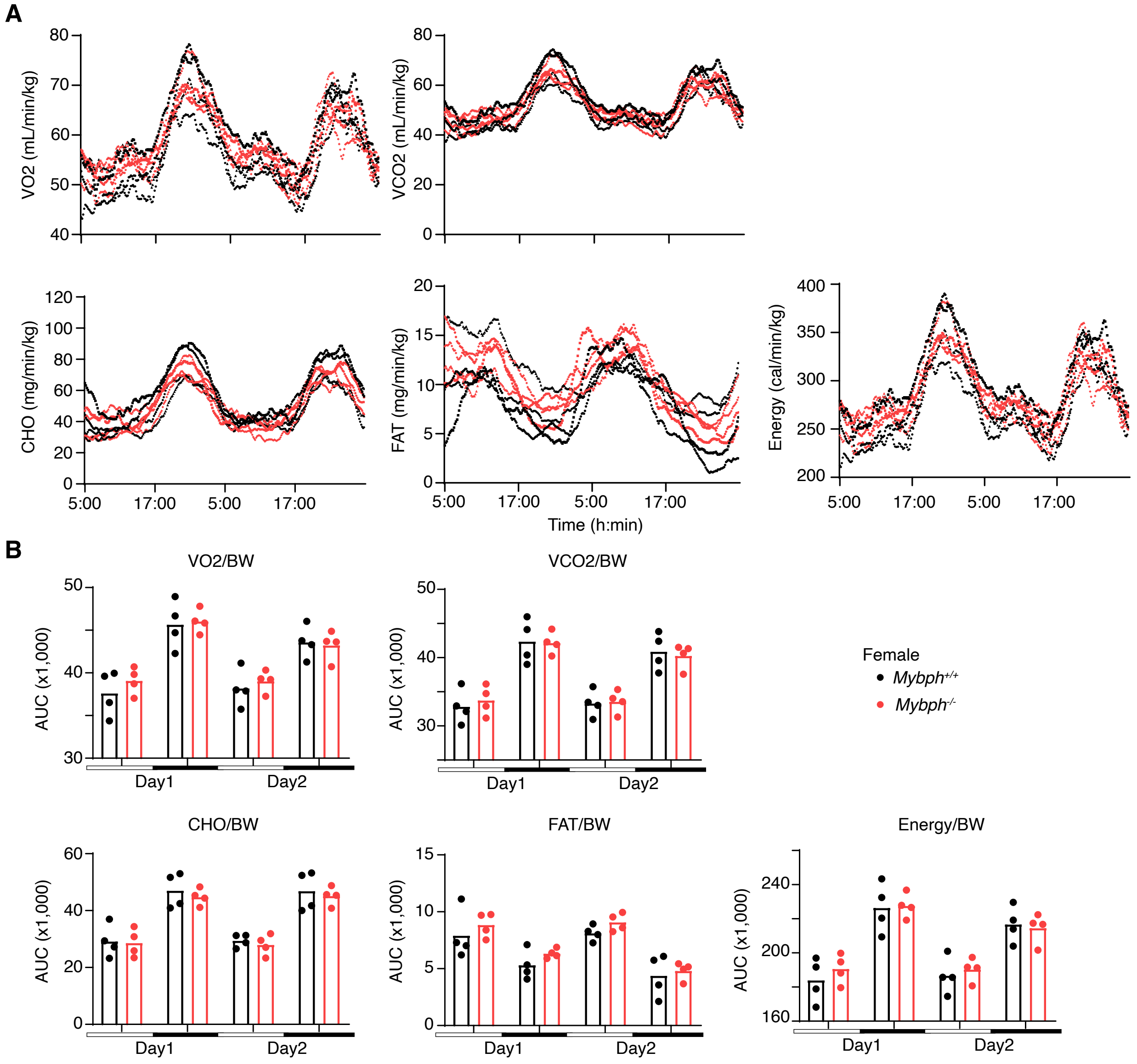


**Fig. S5. Metabolic assessment of female *Mybph* knockout mice.**
(A) Measurement of oxygen consumption (VO2), carbon dioxide production (VCO2), carbohydrate oxidation (CHO), fat oxidation (FAT), and energy expenditure in female control and *Mybph* knockout mice (n = 4 mice per genotype). Representative data from two consecutive days are shown. (B) Quantification of VO2, VCO2, CHO, FAT, and energy expenditure normalized to body weight (BW) (n = 4 mice per genotype). White and black bars indicate daytime and nighttime, respectively.

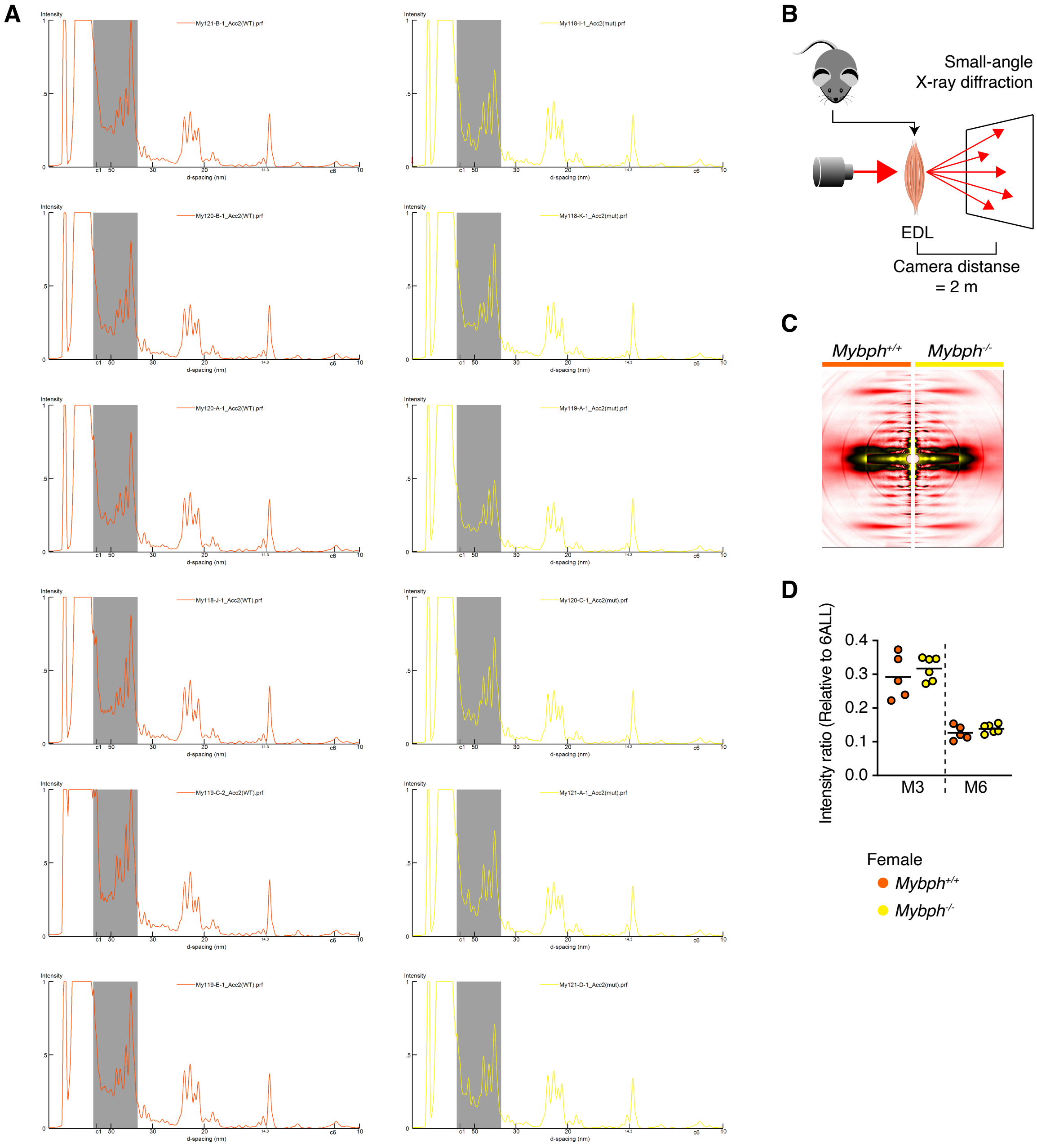


**Fig. S6. Small-angle X-ray scattering from female *Mybph* knockout mice.**
(A) Individual one-dimensional intensity profiles from EDL muscles of female control and *Mybph* knockout mice (n = 6 mice per genotype). Areas under the curves within the shaded gray region (d = 35 to 70 nm) were calculated and normalized to the M3 intensity. (B) Schematic of small-angle X-ray scattering analysis of extensor digitorum longus (EDL) muscles at a sample-to-detector distance of 2 m. (C) Representative X-ray scattering patterns from EDL muscles of female control and *Mybph* knockout mice. (D) Integrated intensities of the third-order myosin meridional reflection (d = 14.3 nm, M3) and sixth-order myosin meridional reflection (d = 7.15 nm, M6), normalized to the actin sixth layer-line reflection (ALL6), whose integrated intensity was set to 1.

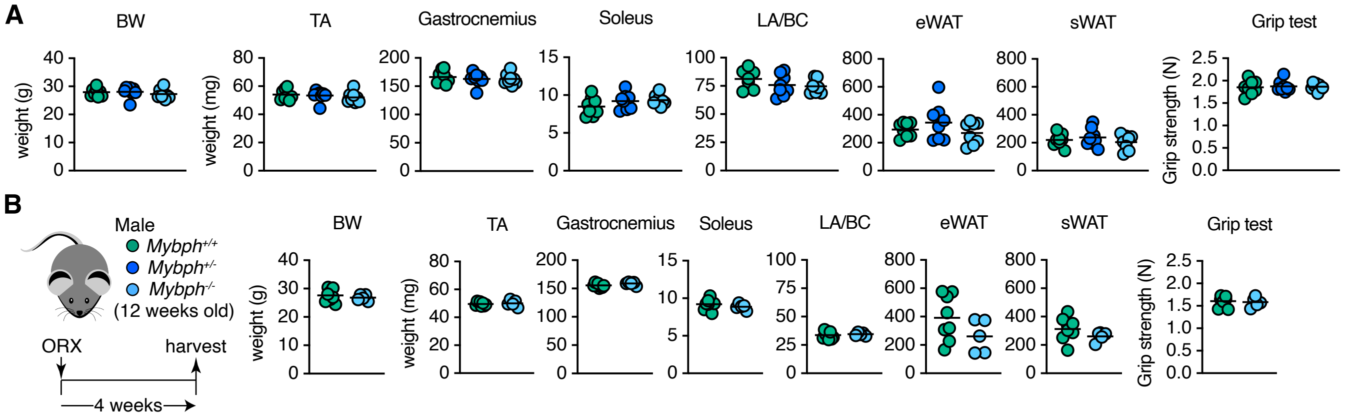


**Fig. S7. Analysis of male *Mybph* knockout mice.**
(A) Quantification of body weight (BW), muscle mass, eWAT and sWAT mass, and grip strength in 12-week-old male control, *Mybph* heterozygous, and homozygous knockout mice (n = 8 mice per genotype). TA, tibialis anterior; LA, levator ani; BC, bulbocavernosus; WAT, white adipose tissue. (B) Experimental design for orchiectomy (ORX) followed by sample collection in male control and *Mybph* knockout mice (left), and quantification of BW, muscle mass, WAT mass, and grip strength (right; n = 5 mice per genotype).

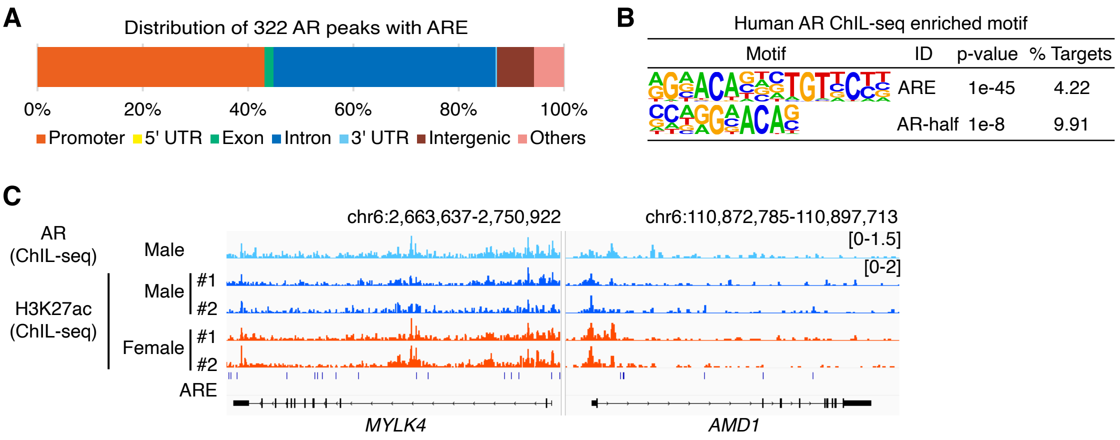


**Fig. S8. Analysis of a human skeletal muscle AR ChIL-seq data set.**
(A) Genomic distribution of ARE-containing AR peaks detected by ChIL-seq in human skeletal muscle. (B) Enriched motifs at AR peaks detected by ChIL-seq in human skeletal muscle. (C) AR binding and histone modifications detected by ChIL-seq in human skeletal muscle at androgen- and AR-regulated genes identified in mouse skeletal muscle.

**Table S1. Primer and guide RNA (gRNA) sequences used in this study.**

| Name | Sequence 5’ to 3’ | Application |
|---|---|---|
| AirID-F | ATGAAGGACAACACCGTTCC | Genotyping |
| AirID-R | AGCAGATCTGAGGCTGATCTC | Genotyping |
| Mybph-F1 | CACAGCTGGGAATGGTTTG | Genotyping |
| Mybph-R1 | CCCAGCTCACAGTCAAGGA | Genotyping |
| AirID-AR | TCAAGGATGGAGGTGCAGTT | gRNA for $Ar^{AirID/Y}$ mice |
| Mybph #1 | TGGAGCCTGCTCTTGCCCGG | gRNA for $Mybph^{-/-}$ mice |
| Mybph #2 | AGGCTTAGTGGCTGCGGGCA | gRNA for $Mybph^{-/-}$ mice |

**Data S1. (separate file)**
Raw data of detected peptides from TA muscles of AirID-AR mice without treatment.

**Data S2. (separate file)**
Analyzed data of detected peptides from TA muscles of AirID-AR mice without treatment.

**Data S3. (separate file)**
Raw data of detected peptides from TA muscles of AirID-AR mice treated with DHT.

**Data S4. (separate file)**
Analyzed data of detected peptides from TA muscles of AirID-AR mice treated with DHT.
